# Comparative efficacy and acceptability of non-surgical brain stimulation for the acute treatment of adult major depressive episodes: A systematic review and network meta-analysis of 113 randomised clinical trials

**DOI:** 10.1101/426866

**Authors:** Julian Mutz, Vijeinika Vipulananthan, Ben Carter, René Hurlemann, Cynthia HY Fu, Allan H Young

**Affiliations:** Social Genetic and Developmental Psychiatry Centre, Institute of Psychiatry, Psychology and Neuroscience, King’s College London, London, United Kingdom; South London and Maudsley Foundation NHS Trust, London, United Kingdom; Department of Biostatistics, Institute of Psychiatry, Psychology and Neuroscience, King’s College London, London, United Kingdom; Department of Psychiatry & Division of Medical Psychology, University of Bonn Medical Centre, Bonn, Germany; School of Psychology, College of Applied Health and Communities, University of East London, London, United Kingdom; Centre for Affective Disorders, Department of Psychological Medicine, Institute of Psychiatry, Psychology and Neuroscience, King’s College London, London, United Kingdom

**Author notes:** Corresponding author: Mr Julian Mutz, Social Genetic and Developmental Psychiatry Centre, Institute of Psychiatry, Psychology and Neuroscience, King’s College London, Memory Lane, London SE5 8AF, United Kingdom.

## Abstract

**Background:** Non-surgical brain stimulation techniques have been applied as tertiary treatments in major depression. However, the relative efficacy and acceptability of individual protocols is uncertain. Our aim was to estimate the comparative clinical efficacy and acceptability of non-surgical brain stimulation for the acute treatment of major depressive episodes in adults.

**Methods:** Embase, PubMed/MEDLINE and PsycINFO were searched up until May 8, 2018, supplemented by manual searches of bibliographies of recent reviews and included trials. We included clinical trials with random allocation to electroconvulsive therapy (ECT), repetitive transcranial magnetic stimulation (rTMS), accelerated TMS (aTMS), priming TMS (pTMS), deep TMS (dTMS), theta burst stimulation (TBS), synchronised TMS (sTMS), magnetic seizure therapy (MST) or transcranial direct current stimulation (tDCS) protocols or sham. Data were extracted from published reports and outcomes were synthesised using pairwise and network random-effects meta-analysis. Primary outcomes were response (efficacy) and all-cause discontinuation (acceptability). We computed odds ratios (OR) with 95% confidence intervals (CI). Remission and continuous post-treatment depression severity scores were also examined.

**Results:** 113 trials (262 treatment arms) randomising 6,750 patients (mean age = 47.9 years; 59% female) with major depressive disorder or bipolar depression met our inclusion criteria. In terms of efficacy, 10 out of 18 treatment protocols were associated with higher response relative to sham in network meta-analysis: bitemporal ECT (OR=8.91, 95%CI 2.57-30.91), high-dose right-unilateral ECT (OR=7.27, 1.90-27.78), pTMS (OR=6.02, 2.21-16.38), MST (OR=5.55, 1.06-28.99), bilateral rTMS (OR=4.92, 2.93-8.25), bilateral TBS (OR=4.44, 1.47-13.41), low-frequency right rTMS (OR=3.65, 2.13-6.24), intermittent TBS (OR=3.20, 1.45-7.08), high-frequency left rTMS (OR=3.17, 2.29-4.37) and tDCS (OR=2.65, 1.55-4.55). Comparing active treatments, bitemporal ECT and high-dose right-unilateral ECT were associated with increased response. All treatment protocols were at least as acceptable as sham treatment.

**Conclusion:** We found that non-surgical brain stimulation techniques constitute viable alternative or add-on treatments for adult patients with major depressive episodes. Our findings also highlight the need to consider other patient and treatment-related factors in addition to antidepressant efficacy and acceptability when making clinical decisions; and emphasize important research priorities in the field of brain stimulation.

Treatment abbreviations
ECT = Electroconvulsive Therapy

- BF ECT = bifrontal ECT **(1)**
- BT ECT = bitemporal ECT **(2)**
- RUL ECT= right unilateral ECT

- H-RUL = high-dose RUL ECT **(3)**
- LM-RUL = low to moderate-dose RUL ECT **(4)**
rTMS = repetitive Transcranial Magnetic Stimulation

- HF-L rTMS = high-frequency rTMS of the left DLPFC **(5)**
- HF-R rTMS = high-frequency rTMS of the right DLPFC **(6)**
- LF-R rTMS = low-frequency rTMS of the right DLPFC **(7)**
- LF-L rTMS = low-frequency rTMS of the left DLPFC **(8)**
- BL rTMS = bilateral rTMS of the DLPFC **(9)**
dTMS = deep Transcranial Magnetic Stimulation **(10)**
pTMS = priming Transcranial Magnetic Stimulation **(11)**
aTMS = accelerated Transcranial Magnetic Stimulation **(12)**
sTMS = synchronised Transcranial Magnetic Stimulation **(13)**
TBS = Theta Burst Stimulation

- iTBS = intermittent TBS of the left DLPFC **(14)**
- cTBS = continuous TBS of the right DLPFC **(15)**
- blTBS = bilateral TBS of the DLPFC **(16)**
MST = Magnetic Seizure Therapy **(17)**
tDCS = transcranial Direct Current Stimulation **(18)**

**Key points**

**Question:** What is the clinical efficacy and acceptability of non-surgical brain stimulation protocols for the acute treatment of major depressive episodes in adults?

**Findings:** In this network meta-analysis, 10 out of 18 treatment protocols were associated with higher response rates relative to sham, most notably bitemporal and high-dose right unilateral electroconvulsive therapy. All treatment protocols were at least as acceptable as sham treatment.

**Meaning:** Non-surgical brain stimulation techniques constitute viable alternative or add-on treatment strategies for adult patients with major depressive episodes.

## Background

Major depression is a highly prevalent and debilitating illness^1^ with considerable disease burden^2^. Its disease course is often recurrent and can become chronic, with relapse rates of up to 80% within one year of remission^3^. Multiple treatment strategies are available - pharmacological interventions and psychological therapies are the most frequently prescribed treatments. However, the effectiveness of these treatments remains limited and less than 50% of patients respond to an initial course of drug treatment^4^. A significant number of patients do not tolerate pharmacotherapy because of undesired effects including sexual dysfunction, weight gain and insomnia^5,6^. Combination strategies with multiple pharmacological agents increase the risk for adverse events and drug interactions^7^. These factors limit medication-adherence and potentially cause discontinuation of treatment^8^. Similarly, psychological therapies are not effective for every patient and may also be associated with undesired effects^9^.

Non-surgical brain stimulation techniques including electroconvulsive therapy (ECT) and repetitive transcranial magnetic stimulation (rTMS) have been applied as tertiary treatments or are considered to be alternative or add-on treatments for major depressive episodes. Over the past decade, novel modifications of standard rTMS have been developed to optimize treatment: deep transcranial magnetic stimulation (dTMS), theta burst stimulation (TBS), priming transcranial magnetic stimulation (pTMS), accelerated transcranial magnetic stimulation (aTMS) and synchronised transcranial magnetic stimulation (sTMS). Clinical trials have also examined the antidepressant efficacy of magnetic seizure therapy (MST) and transcranial direct current stimulation (tDCS) (Supplement 1).

Previous meta-analyses have examined the clinical efficacy and acceptability of brain stimulation compared to placebo^10^ or within pairs of active treatments^11^. However, these approaches provide limited insights into the overall treatment hierarchy because the available evidence was not synthesised in one step. Moreover, the absence of head-to-head clinical trials for some treatment comparisons creates uncertainty for decision-makers.

Network meta-analysis (NMA) includes both direct and indirect treatment comparisons^12^, and should be regarded as the highest level of evidence in treatment guidelines^13^ and may overcome a lack of head to head evaluation. Two NMAs of brain stimulation therapies for major depressive episodes have been published but were limited in scope of included interventions^14,15^. The NMA by Brunoni et al.^14^ provided a comprehensive synthesis of the available evidence for rTMS, but did not include ECT, MST or tDCS. Moreover, studies that had co-initiated pharmacotherapy were included in their analyses, potentially inflating efficacy estimates of rTMS. The Chen et al.^15^ NMA included trials that had compared rTMS to ECT, but did not include sham-controlled trials or distinguish the various electrode placements or electrical dosages of ECT.

### Objective

The primary aim of this study is to estimate the efficacy and acceptability of non-surgical brain stimulation protocols for the acute treatment of major depressive episodes in adults participating in randomised clinical trials (RCTs).

## Methods

We followed the PRISMA guidelines for NMA^16^. The study was conducted between January 17, 2017 and September 14, 2018. No review protocol or registration are available.

### Criteria for considering studies for this review

We included RCTs with parallel-group or cross-over designs. Only data from period one were analysed to avoid potential carry-over effects. Studies needed to include a clinician-administered depression rating scale, the Hamilton Depression Rating Scale (HDRS)^17^ or the Montgomery-Åsberg Depression Rating Scale (MADRS)^18^. Conference abstracts, editorials, reviews, meta-analyses and case reports or case series were excluded. We also excluded non-English language publications and those reporting duplicate data.

Participants had to be adults (age ≥ 18 years) with RDC, DSM or ICD diagnosis of major depressive disorder (MDD) or bipolar depression. Other primary diagnoses were excluded, as were trials that recruited patients with a subtype of depression (e.g. postpartum depression) or with depression as secondary diagnosis (e.g. fibromyalgia and depression). Finally, we excluded non-human studies.

Studies had to include at least two of the following treatments: tDCS, rTMS, TBS, dTMS, sTMS, pTMS, aTMS, ECT, MST or sham. For rTMS, protocols were grouped according to coil location and stimulation frequency: high-frequency stimulation of the left dorsolateral prefrontal cortex (DLPFC; HF-L), high-frequency stimulation of the right DLPFC (HF-R), low-frequency stimulation of the right DLPFC (LF-R), low-frequency stimulation of the left DLPFC (LF-L) and bilateral stimulation of the DLPFC (BL). TBS protocols were grouped in a similar fashion: intermittent stimulation of the left DLPFC (iTBS), continuous stimulation of the right DLPFC (cTBS) and bilateral stimulation of the DLPFC (blTBS). Finally, ECT protocols were grouped according to electrode placement (BF = bifrontal; BT = bitemporal; RUL = right unilateral), and for RUL ECT also according to electrical dosage (H-RUL = high-dose right unilateral; LM-RUL = low to moderate-dose right unilateral). For multi-arm trials, treatment groups that could not be included individually were combined^19^. All sham controls were merged into one node. Supplement 2 shows the network of potential treatment comparisons. We assume that any patient enrolled in one of the trials included in our review is, in principle, equally likely to be randomised to any other trial in the network.

Studies examining vagus nerve stimulation or related interventions were excluded. We also excluded trials in which pharmacological or psychological treatments were co-initiated with brain stimulation.

### Search methods for identification of studies

The Embase, PubMed/MEDLINE and PsycINFO databases (accessed via Ovid) were searched for articles published between the first date available and May 8th, 2018. A full description of our search methods can be found in Supplement 3. Two authors (JM & VV) independently performed the literature search, screened titles and abstracts, selected relevant full-texts and assessed these for eligibility.

### Data extraction

One author (JM) extracted relevant information from eligible trials and a second author (VV) independently reviewed these data. Discrepancies were resolved by consensus. Data that could not be retrieved from the original publications were requested from the corresponding authors or searched for in other reviews. We used WebPlotDigitizer (https://apps.automeris.io/wpd/) to extract numerical data from figures.

### Participant characteristics

Sex (*n* male/female); age in years (mean, standard deviation and range); hospitalisation status (outpatient, inpatient or mixed); whether patients with psychotic symptoms were excluded from the trial (yes/no); diagnosis (MDD, bipolar depression or mixed); treatment strategy (monotherapy, add-on therapy or mixed); and whether patients were considered treatment resistant (yes, no or mixed).

### Intervention characteristics

ECT: electrical dosage (multiples of seizure threshold) and electrode placement. rTMS: coil location and stimulation frequency (in hertz). Similar data were extracted for TBS, also including the treatment protocol (iTBS, cTBS or blTBS).

### Study design and outcomes

Cross-over design (yes/no); HDRS version; response and remission criteria; *n* patients randomised; *n* patients meeting response and remission criteria at primary treatment endpoint; *n* patients discontinuing treatment for any reason; and *n* patients analysed.

### Risk of bias assessment

The Cochrane tool for assessing risk of bias in randomised trials^20^ was used to evaluate each study. Potential sources of bias include random sequence generation, allocation concealment, blinding of participants and personnel, blinding of outcome assessment, incomplete outcome data and selective reporting. Each trial received a study-level score of low, high or unclear risk of bias for each domain. Two authors (JM & VV) independently conducted this assessment and discrepancies were resolved by consensus.

### Data synthesis

We computed odds ratios (ORs; Mantel-Haenszel method) and standardised mean differences (SMD; Hedge’s g) with 95% confidence intervals (CIs) to estimate effect sizes for categorical and continuous outcomes, respectively. The primary outcome measure of efficacy was response, defined in most trials as a ≥ 50% reduction in depressive symptoms at primary treatment endpoint. Remission was our secondary outcome measure of efficacy, according to the criteria used in each trial (e.g. HDRS ≤ 7 at primary treatment endpoint). Continuous post-treatment depression severity scores constituted our tertiary efficacy outcome measure. If trials reported data on both HDRS and MADRS, the HDRS data were selected for analyses to facilitate comparability between trials. In case of multiple HDRS versions, the original 17-item version was analysed. Data based on the intention-to-treat (ITT) or modified intention to treat (mITT) sample were preferred over data based on completers for all analyses.

### Pairwise meta-analysis

We conducted frequentist random-effects meta-analyses of all direct treatment comparisons, allowing for heterogeneity in treatment effects between studies. All pairwise analyses were conducted using the ‘meta’ package^21^ in RStudio 1.0.143.

Statistical heterogeneity within each pairwise comparison was estimated using the *I^2^* statistic, with values of 25%, 50% and 75% representing little, substantial and severe level of heterogeneity ^22^. When severe heterogeneity was exhibited, this was investigated using subgroups to explore the effect modifiers. Subgroups included treatment resistance, diagnosis, hospitalisation status and exclusion of patients with psychotic features.

### Network meta-analysis

Network plots were produced for each outcome to visualise network geometry and node connectivity^23^. NMAs were fit within a frequentist framework using a multivariate random-effects meta-analysis model^24,25^ that accounts for the correlations between effect sizes in trials with more than two groups.

We assumed network consistency and a common heterogeneity parameter across all treatment contrasts. Relative ORs or SMDs and 95% CIs for all treatment comparisons were presented in league tables. We also present relative treatment effects with 95% CIs and 95% prediction intervals (PrIs) for all sham comparisons in forest plots. To obtain treatment hierarchies, we computed ranking probabilities for all ranks and outcomes using a parametric bootstrap procedure with 10,000 resamples^25^. All NMAs were conducted using the ‘mvmeta’^26,27^ and ‘network’^28^ packages in Stata SE 15.0.

We assessed the transitivity assumption by comparing the distribution/frequency of potential effect modifiers across treatment comparisons: continuous (depression severity at baseline, age, percent female) and categorical (treatment resistance, diagnosis, hospitalisation status, exclusion of patients with psychotic features and treatment strategy).

Assuming equivalence of direct and indirect evidence (i.e. consistency) in NMA may lead to inaccurate conclusions when there is evidence for significant inconsistency^25^. We assessed the assumption of consistency by fitting a design-by-treatment interaction model^24,25^ that accounts for loop and design inconsistencies and provides a global Wald test to evaluate inconsistency in the entire network.

We also computed inconsistency factors (IFs) and 95% CIs for each closed triangular and quadratic loop within treatment networks to estimate absolute differences between direct and indirect evidence. We used a method of moments estimator of loop-specific heterogeneity, assuming a common heterogeneity parameter for all comparisons within the same loop.

### Sensitivity analysis

We conducted two sensitivity analyses to assess the robustness of our findings for response and all-cause discontinuation rates: (1) trials that examined tDCS were excluded and (2) trials with high overall risk of bias were excluded.

## Results

The PRISMA flowchart is presented in Figure 1. 113 RCTs (262 treatment arms) met our inclusion criteria. Full citations can be found in Supplement 4-5.

**Figure 1.**
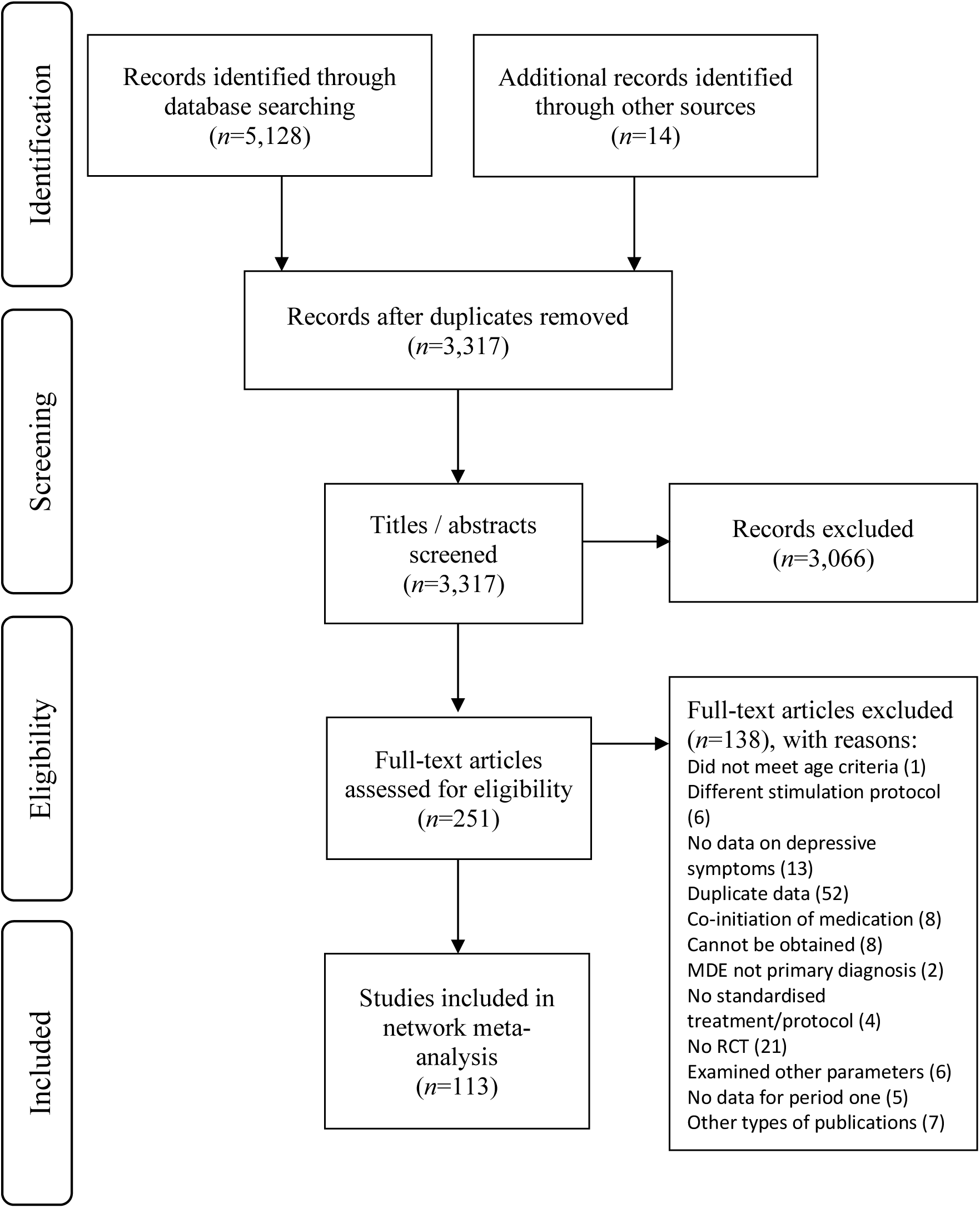
PRISMA flow diagram.

Overall, *N*=6,750 patients were randomised to treatment. The mean age was 47.9 years and 59% (*n* = 3,545) were women. The median study sample size was 40 patients (range=6-414). The risk of bias assessment is presented in Supplement 6. Briefly, 23.9% of the included trials were considered low risk, while 57.5% and 18.6% were categorised as unclear or high risk, respectively.

Most trials (80.9%) recruited only patients with treatment resistant depression (TRD), typically defined as a minimum of two failed pharmacological treatments. Only 12.8% recruited both TRD and non-TRD patients; the remaining 6.4% recruited patients with non-TRD. 58.5% of the studies excluded patients with psychotic features. 49.1% recruited patients with MDD only. For the trials that recruited both patients with MDD and bipolar depression (46.2%), few patients were diagnosed with bipolar depression. 48.8% of trials recruited outpatients only, whereas 29.1% and 22.1% recruited inpatients only or both outpatients and inpatients, respectively. In 63.2% of the studies brain stimulation was an add-on treatment to stable pharmacotherapy in most, if not all, patients. Baseline depression severity, percent female and age were similar across most treatment comparisons. As such, the assumption of transivity is likely to hold in our data.

### Pairwise meta-analysis

The results of the pairwise meta-analysis and heterogeneity estimates are presented in Supplement 7. Briefly, BT ECT, HF-L rTMS, LF-R rTMS, tDCS and dTMS were more efficacious than sham across all outcomes (ORs=1.69 [min] to 5.50 [max]; SMDs=-0.29 [min] to -0.77 [max]). BL rTMS was more efficacious than sham when considering response (4.93, 2.78-8.75; *I*^2^=0%) and remission (4.67, 1.84-11.84; *I*^2^=0%), while iTBS was more efficacious than sham in terms of response (4.25, 1.22-14.84; *I*^2^=0%). There were few differences between active treatments. Most notably, BT ECT was more efficacious than LM-RUL ECT across all outcomes (ORs=3.87 [min] to 6.67 [max]; SMD=-0.88, -1.28 to -0.49; *I*^2^=0%). In terms of all-cause discontinuation, we found no differences between active treatments and sham and heterogeneity between trials could not be explained by stratifying analyses according to hospitalisation status, psychotic symptoms, treatment resistance or diagnosis.

### Network meta-analysis

The results of the NMA of the primary outcome of efficacy (response) and acceptability (all-cause discontinuation) are presented in Table 1.

**Table 1.** Network meta-analysis of response and all-cause discontinuation rates.

|  | tDCS | sTMS | pTMS | iTBS | dTMS | cTBS | biTBS | aTMS | MST | LMRUL | LFR | LFL | HRUL | HFR | HFL | BT | BL | BF | SHM |
| --- | --- | --- | --- | --- | --- | --- | --- | --- | --- | --- | --- | --- | --- | --- | --- | --- | --- | --- | --- |
| tDCS |  | 1.62<br>(0.70,3.78) | **5.46**<br>**(1.63,18.31)** | 1.17<br>(0.47,2.91) | 1.37<br>(0.59,3.20) | 2.55<br>(0.23,28.01) | 3.12<br>(0.31,31.59) | 0.86<br>(0.19,3.83) | 0.69<br>(0.12,4.03) | 1.48<br>(0.54,4.06) | 1.28<br>(0.56,2.92) | 0.58<br>(0.14,2.41) | 1.47<br>(0.59,3.66) | 1.30<br>(0.04,42.23) | 1.41<br>(0.80,2.51) | 1.56<br>(0.70,3.45) | **2.38**<br>**(1.15,4.94)** | 1.50<br>(0.59,3.82) | 1.17<br>(0.72,1.91) |
| sTMS | 1.27<br>(0.41,3.95) |  | 3.37<br>(0.92,12.38) | 0.72<br>(0.26,2.03) | 0.85<br>(0.32,2.24) | 1.57<br>(0.14,18.13) | 1.93<br>(0.18,20.47) | 0.53<br>(0.11,2.55) | 0.43<br>(0.07,2.65) | 0.91<br>(0.30,2.79) | 0.79<br>(0.30,2.05) | 0.36<br>(0.08,1.61) | 0.91<br>(0.32,2.54) | 0.80<br>(0.02,26.93) | 0.87<br>(0.41,1.84) | 0.96<br>(0.38,2.44) | 1.47<br>(0.61,3.52) | 0.92<br>(0.32,2.65) | 0.72<br>(0.36,1.44) |
| pTMS | 0.44<br>(0.14,1.36) | 0.35<br>(0.08,1.43) |  | **0.21**<br>**(0.06,0.81)** | **0.25**<br>**(0.07,0.92)** | 0.47<br>(0.04,6.19) | 0.57<br>(0.05,6.85) | **0.16**<br>**(0.03,0.93)** | **0.13**<br>**(0.02,0.95)** | 0.27<br>(0.07,1.11) | **0.23**<br>**(0.08,0.72)** | **0.11**<br>**(0.02,0.59)** | 0.27<br>(0.07,1.02) | 0.24<br>(0.01,8.84) | **0.26**<br>**(0.08,0.79)** | 0.28<br>(0.08,1.01) | 0.44<br>(0.16,1.16) | 0.27<br>(0.07,1.06) | **0.21**<br>**(0.07,0.65)** |
| iTBS | 0.83<br>(0.32,2.16) | 0.65<br>(0.18,2.37) | 1.88<br>(0.54,6.57) |  | 1.17<br>(0.42,3.28) | 2.17<br>(0.19,24.42) | 2.66<br>(0.26,27.58) | 0.73<br>(0.15,3.49) | 0.59<br>(0.09,3.71) | 1.26<br>(0.40,3.95) | 1.09<br>(0.41,2.92) | 0.49<br>(0.11,2.27) | 1.25<br>(0.44,3.58) | 1.10<br>(0.03,37.38) | 1.20<br>(0.59,2.47) | 1.33<br>(0.51,3.44) | 2.03<br>(0.82,5.05) | 1.28<br>(0.44,3.73) | 1.00<br>(0.46,2.14) |
| dTMS | 1.42<br>(0.51,3.95) | 1.12<br>(0.29,4.25) | 3.22<br>(0.85,12.11) | 1.71<br>(0.53,5.58) |  | 1.86<br>(0.16,21.41) | 2.27<br>(0.21,24.18) | 0.63<br>(0.13,3.01) | 0.50<br>(0.08,3.13) | 1.08<br>(0.35,3.29) | 0.93<br>(0.36,2.42) | 0.42<br>(0.09,1.90) | 1.07<br>(0.38,3.01) | 0.94<br>(0.03,31.80) | 1.03<br>(0.49,2.18) | 1.13<br>(0.45,2.88) | 1.74<br>(0.72,4.16) | 1.09<br>(0.38,3.13) | 0.85<br>(0.43,1.70) |
| cTBS | 2.58<br>(0.66,10.06) | 2.03<br>(0.41,10.18) | **5.84**<br>**(1.19,28.61)** | 3.11<br>(0.80,12.08) | 1.82<br>(0.39,8.37) |  | 1.23<br>(0.06,24.03) | 0.34<br>(0.02,5.18) | 0.27<br>(0.02,4.89) | 0.58<br>(0.05,7.11) | 0.50<br>(0.04,5.72) | 0.23<br>(0.02,3.38) | 0.58<br>(0.05,6.79) | 0.51<br>(0.01,32.89) | 0.55<br>(0.05,5.86) | 0.61<br>(0.05,6.91) | 0.93<br>(0.08,10.30) | 0.59<br>(0.05,6.98) | 0.46<br>(0.04,4.80) |
| biTBS | 0.60<br>(0.17,2.04) | 0.47<br>(0.11,2.11) | 1.35<br>(0.32,5.71) | 0.72<br>(0.21,2.45) | 0.42<br>(0.10,1.72) | 0.23<br>(0.05,1.02) |  | 0.27<br>(0.02,3.93) | 0.22<br>(0.01,3.72) | 0.47<br>(0.04,5.35) | 0.41<br>(0.04,4.23) | 0.19<br>(0.01,2.56) | 0.47<br>(0.04,5.11) | 0.42<br>(0.01,25.60) | 0.45<br>(0.05,4.39) | 0.50<br>(0.05,5.19) | 0.76<br>(0.08,7.52) | 0.48<br>(0.04,5.25) | 0.37<br>(0.04,3.60) |
| aTMS | 1.31<br>(0.36,4.69) | 1.03<br>(0.22,4.80) | 2.96<br>(0.66,13.40) | 1.58<br>(0.40,6.20) | 0.92<br>(0.22,3.94) | 0.51<br>(0.09,2.77) | 2.19<br>(0.45,10.71) |  | 0.81<br>(0.09,7.19) | 1.73<br>(0.33,8.96) | 1.49<br>(0.32,6.98) | 0.67<br>(0.10,4.67) | 1.71<br>(0.35,8.36) | 1.51<br>(0.04,62.09) | 1.65<br>(0.41,6.60) | 1.81<br>(0.40,8.32) | 2.78<br>(0.62,12.41) | 1.74<br>(0.35,8.64) | 1.36<br>(0.33,5.60) |
| MST | 0.48<br>(0.08,2.72) | 0.38<br>(0.05,2.63) | 1.08<br>(0.16,7.41) | 0.58<br>(0.09,3.54) | 0.34<br>(0.05,2.19) | 0.19<br>(0.02,1.47) | 0.80<br>(0.11,5.81) | 0.37<br>(0.05,2.69) |  | 2.14<br>(0.42,10.78) | 1.85<br>(0.30,11.27) | 0.84<br>(0.10,7.18) | 2.12<br>(0.44,10.26) | 1.87<br>(0.04,86.69) | 2.04<br>(0.37,11.10) | 2.25<br>(0.44,11.37) | 3.44<br>(0.59,20.15) | 2.16<br>(0.43,10.88) | 1.69<br>(0.31,9.16) |
| LMRUL | 0.97<br>(0.25,3.68) | 0.76<br>(0.16,3.74) | 2.19<br>(0.46,10.50) | 1.17<br>(0.28,4.88) | 0.68<br>(0.15,3.07) | 0.38<br>(0.07,2.14) | 1.62<br>(0.31,8.33) | 0.74<br>(0.14,3.87) | 2.02<br>(0.64,6.36) |  | 0.86<br>(0.29,2.57) | 0.39<br>(0.08,1.93) | 0.99<br>(0.43,2.31) | 0.87<br>(0.02,30.58) | 0.95<br>(0.39,2.33) | 1.05<br>(0.47,2.33) | 1.61<br>(0.58,4.47) | 1.01<br>(0.46,2.22) | 0.79<br>(0.33,1.90) |
| LFR | 0.73<br>(0.34,1.55) | 0.57<br>(0.18,1.80) | 1.65<br>(0.62,4.42) | 0.88<br>(0.35,2.21) | 0.51<br>(0.18,1.43) | 0.28<br>(0.07,1.09) | 1.22<br>(0.37,4.03) | 0.56<br>(0.16,1.94) | 1.52<br>(0.27,8.51) | 0.75<br>(0.20,2.79) |  | 0.45<br>(0.11,1.95) | 1.15<br>(0.42,3.13) | 1.01<br>(0.03,33.81) | 1.10<br>(0.56,2.19) | 1.22<br>(0.50,2.99) | 1.86<br>(0.99,3.49) | 1.17<br>(0.42,3.26) | 0.91<br>(0.47,1.77) |
| LFL | 2.39<br>(0.42,13.77) | 1.89<br>(0.27,13.26) | 5.42<br>(0.79,37.15) | 2.89<br>(0.46,17.99) | 1.69<br>(0.26,11.08) | 0.93<br>(0.12,7.43) | 4.00<br>(0.55,29.34) | 1.83<br>(0.24,13.67) | 5.00<br>(0.48,51.70) | 2.47<br>(0.32,19.25) | 3.29<br>(0.59,18.32) |  | 2.54<br>(0.55,11.82) | 2.24<br>(0.06,90.30) | 2.44<br>(0.63,9.45) | 2.69<br>(0.62,11.72) | 4.12<br>(0.99,17.17) | 2.59<br>(0.55,12.21) | 2.02<br>(0.53,7.73) |
| HRUL | 0.36<br>(0.09,1.55) | 0.29<br>(0.05,1.55) | 0.83<br>(0.16,4.36) | 0.44<br>(0.09,2.04) | 0.26<br>(0.05,1.28) | **0.14**<br>**(0.02,0.88)** | 0.61<br>(0.11,3.44) | 0.28<br>(0.05,1.60) | 0.76<br>(0.22,2.63) | **0.38**<br>**(0.18,0.78)** | 0.50<br>(0.12,2.08) | 0.15<br>(0.02,1.28) |  | 0.88<br>(0.03,29.99) | 0.96<br>(0.44,2.08) | 1.06<br>(0.61,1.83) | 1.62<br>(0.64,4.09) | 1.02<br>(0.60,1.73) | 0.80<br>(0.37,1.72) |
| HFR | 1.58<br>(0.31,7.97) | 1.24<br>(0.20,7.79) | 3.58<br>(0.58,22.00) | 1.90<br>(0.35,10.48) | 1.11<br>(0.19,6.48) | 0.61<br>(0.09,4.40) | 2.64<br>(0.40,17.31) | 1.21<br>(0.18,8.06) | 3.30<br>(0.35,30.98) | 1.63<br>(0.23,11.38) | 2.17<br>(0.44,10.81) | 0.66<br>(0.07,6.27) | 4.32<br>(0.57,32.63) |  | 1.09<br>(0.03,34.34) | 1.20<br>(0.04,39.79) | 1.84<br>(0.06,60.14) | 1.16<br>(0.03,39.61) | 0.90<br>(0.03,28.44) |
| HFL | 0.84<br>(0.45,1.56) | 0.66<br>(0.23,1.91) | 1.90<br>(0.69,5.22) | 1.01<br>(0.46,2.20) | 0.59<br>(0.23,1.50) | 0.33<br>(0.09,1.17) | 1.40<br>(0.45,4.33) | 0.64<br>(0.21,1.99) | 1.75<br>(0.34,9.11) | 0.87<br>(0.26,2.91) | 1.15<br>(0.67,1.98) | 0.35<br>(0.07,1.86) | 2.30<br>(0.60,8.75) | 0.53<br>(0.11,2.46) |  | 1.10<br>(0.58,2.08) | 1.69<br>(0.95,2.99) | 1.06<br>(0.47,2.37) | 0.83<br>(0.61,1.12) |
| BT | 0.30<br>(0.08,1.15) | 0.23<br>(0.05,1.17) | 0.67<br>(0.14,3.28) | 0.36<br>(0.08,1.53) | **0.21**<br>**(0.05,0.96)** | **0.12**<br>**(0.02,0.67)** | 0.50<br>(0.10,2.61) | 0.23<br>(0.04,1.21) | 0.62<br>(0.18,2.14) | **0.31**<br>**(0.17,0.57)** | 0.41<br>(0.11,1.55) | **0.12**<br>**(0.02,0.98)** | 0.82<br>(0.45,1.49) | 0.19<br>(0.03,1.34) | 0.36<br>(0.10,1.23) |  | 1.53<br>(0.68,3.45) | 0.96<br>(0.53,1.73) | 0.75<br>(0.40,1.41) |
| BL | 0.54<br>(0.26,1.13) | 0.43<br>(0.14,1.32) | 1.22<br>(0.49,3.05) | 0.65<br>(0.26,1.62) | 0.38<br>(0.14,1.05) | <u>0.21</u><br><u>(0.06,0.79)</u> | 0.90<br>(0.29,2.83) | 0.41<br>(0.12,1.44) | 1.13<br>(0.20,6.30) | 0.56<br>(0.15,2.07) | 0.74<br>(0.41,1.35) | 0.23<br>(0.04,1.27) | 1.48<br>(0.36,6.14) | 0.34<br>(0.07,1.70) | 0.64<br>(0.37,1.11) | 1.81<br>(0.48,6.87) |  | 0.63<br>(0.24,1.63) | <u>0.49</u><br><u>(0.29,0.84)</u> |
| BF | 0.78<br>(0.17,3.65) | 0.62<br>(0.11,3.60) | 1.77<br>(0.31,10.14) | 0.94<br>(0.19,4.79) | 0.55<br>(0.10,2.98) | 0.30<br>(0.05,2.04) | 1.31<br>(0.21,7.99) | 0.60<br>(0.10,3.71) | 1.63<br>(0.41,6.44) | 0.81<br>(0.36,1.82) | 1.07<br>(0.23,4.92) | 0.33<br>(0.04,2.93) | 2.14<br>(0.83,5.53) | 0.50<br>(0.06,4.01) | 0.93<br>(0.22,3.93) | <u>2.62</u><br><u>(1.03,6.71)</u> | 1.45<br>(0.32,6.63) |  | 0.78<br>(0.35,1.74) |
| SHM | <u>2.65</u><br><u>(1.55,4.55)</u> | 2.09<br>(0.76,5.77) | <u>6.02</u><br><u>(2.21,16.38)</u> | <u>3.20</u><br><u>(1.45,7.08)</u> | 1.87<br>(0.78,4.49) | 1.03<br>(0.29,3.60) | <u>4.44</u><br><u>(1.47,13.41)</u> | 2.03<br>(0.64,6.48) | <u>5.55</u><br><u>(1.06,28.99)</u> | 2.74<br>(0.81,9.31) | <u>3.65</u><br><u>(2.13,6.24)</u> | 1.11<br>(0.21,5.87) | <u>7.27</u><br><u>(1.90,27.78)</u> | 1.68<br>(0.36,7.77) | <u>3.17</u><br><u>(2.29,4.37)</u> | <u>8.91</u><br><u>(2.57,30.91)</u> | <u>4.92</u><br><u>(2.93,8.25)</u> | 3.40<br>(0.80,14.41) |  |

*Note.* Effect sizes represent relative odds ratios and 95% confidence intervals. For the lower triangle (response rates), values lower than 1 favour the treatment in the corresponding row, while values higher than 1 favour the treatment in the corresponding column. For the upper triangle (all-cause discontinuation rates), values lower than 1 favour the treatment in the corresponding row while values higher than 1 favour the treatment in the corresponding column. SHM = Sham; MST = Magnetic Seizure Therapy; ECT = Electroconvulsive Therapy; LMRUL = Low to Moderate-Dose Right Unilateral ECT; rTMS = repetitive Transcranial Magnetic Stimulation; LFR = Low-Frequency Right rTMS; LFL = Low-Frequency Left rTMS; HRUL = High-Dose Right Unilateral ECT; HFR = High-Frequency Right rTMS; HFL = High-Frequency Left rTMS; BT = Bitemporal ECT; BL = Bilateral rTMS; BF = Bifrontal ECT; tDCS = transcranial Direct Current Stimulation; sTMS = synchronised Transcranial Magnetic Stimulation; pTMS = priming Transcranial Magnetic Stimulation; TBS = Theta Burst Stimulation; iTBS = intermittent TBS; dTMS = deep Transcranial Magnetic Stimulation; cTBS = continuous TBS; bITBS = Bilateral TBS; aTMS = accelerated Transcranial Magnetic Stimulation.

Response rates were available for 208 treatment arms (*n* = 5,962) including all 18 active interventions and sham (Figure 2).

**Figure 2.**
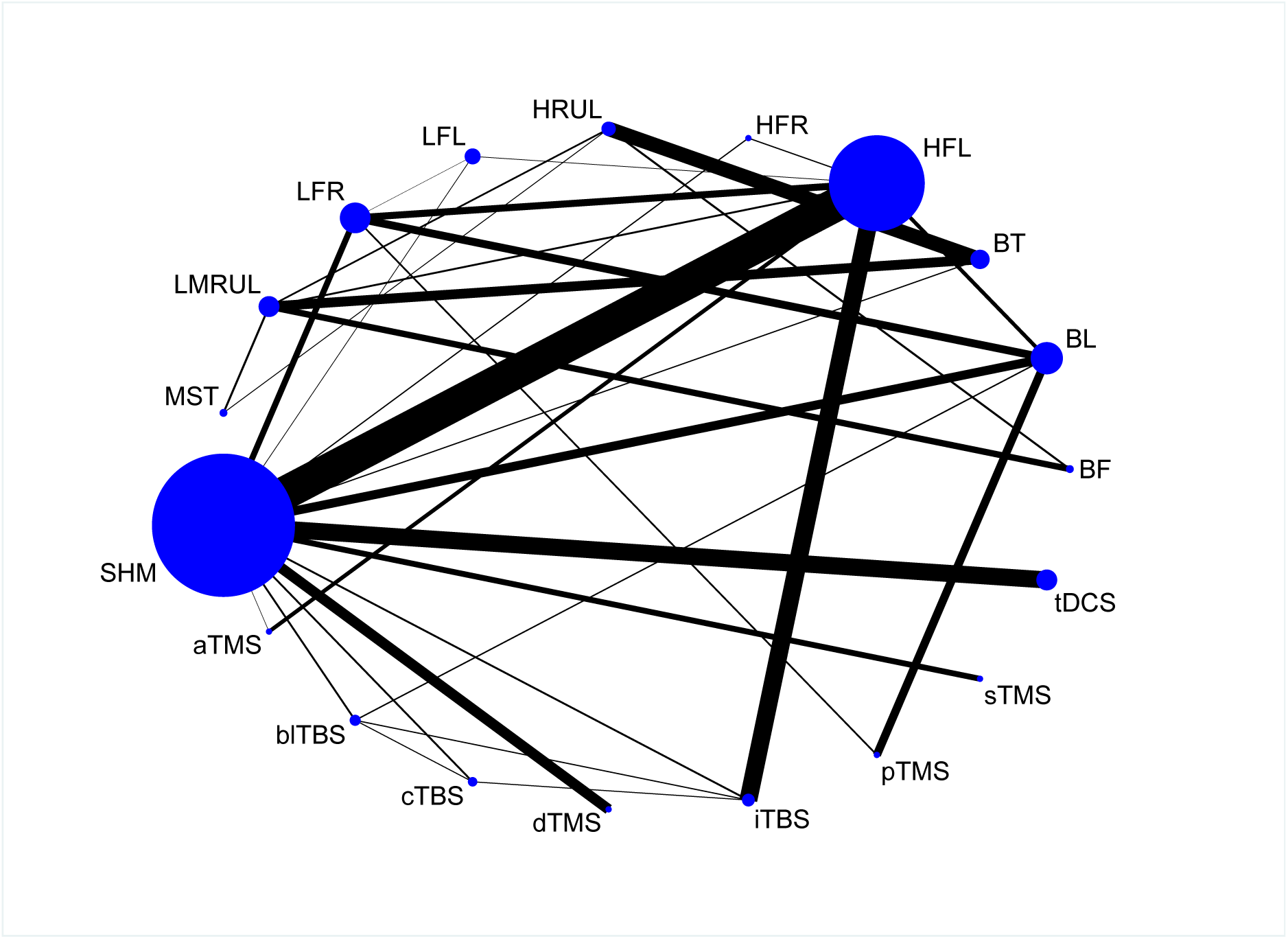
Network plot of available treatment comparisons for response rates. The size of the nodes is proportional to the number of patients randomised to each treatment. The width of the lines is proportional to the number of RCTs comparing each pair of treatments. SHM = Sham; MST = Magnetic Seizure Therapy; ECT = Electroconvulsive Therapy; LMRUL = Low to Moderate-Dose Right Unilateral ECT; rTMS = repetitive Transcranial Magnetic Stimulation; LFR = Low-Frequency Right rTMS; LFL = Low-Frequency Left rTMS; HRUL = High-Dose Right Unilateral ECT; HFR = High-Frequency Right rTMS; HFL = High-Frequency Left rTMS; BT = Bitemporal ECT; BL = Bilateral rTMS; BF = Bifrontal ECT; tDCS = transcranial Direct Current Stimulation; sTMS = synchronised Transcranial Magnetic Stimulation; pTMS = priming Transcranial Magnetic Stimulation; TBS = Theta Burst Stimulation; iTBS = intermittent TBS; dTMS = deep Transcranial Magnetic Stimulation; cTBS = continuous TBS; blTBS = Bilateral TBS; aTMS = accelerated Transcranial Magnetic Stimulation.

The results of the NMA indicate that BT ECT (OR=8.91, 95%CI 2.57-30.91), H-RUL ECT (7.27, 1.90-27.78), pTMS (6.02, 2.21-16.38), MST (5.55, 1.06-28.99), BL rTMS (4.92, 2.93-8.25), blTBS (4.44, 1.47-13.41), LF-R rTMS (3.65, 2.13-6.24), iTBS (3.20, 1.45-7.08), HF-L rTMS (3.17, 2.29-4.37) and tDCS (2.65, 1.55-4.55) were more efficacious than sham (Figure 3). However, MST, iTBS and tDCS did not remain significant when examining prediction intervals.

**Figure 3.**
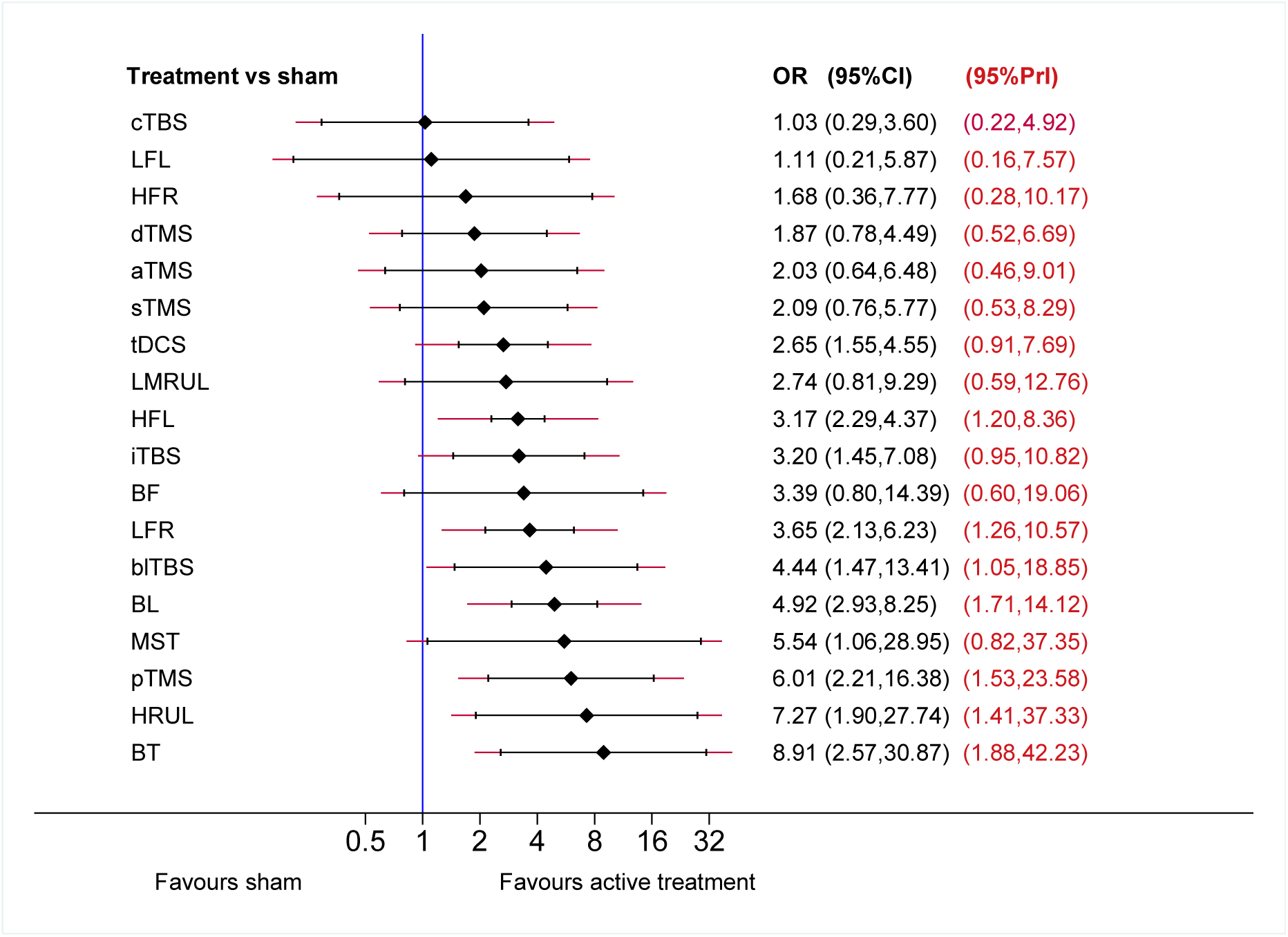
Forest plot of active vs sham treatment comparisons for response rates. Effect sizes represent relative odds ratios (ORs) with 95% confidence intervals (Cis) and 95% prediction intervals (PrIs). SHM = Sham; MST = Magnetic Seizure Therapy; ECT = Electroconvulsive Therapy; LMRUL = Low to Moderate-Dose Right Unilateral ECT; rTMS = repetitive Transcranial Magnetic Stimulation; LFR = Low-Frequency Right rTMS; LFL = Low-Frequency Left rTMS; HRUL = High-Dose Right Unilateral ECT; HFR = High-Frequency Right rTMS; HFL = High-Frequency Left rTMS; BT = Bitemporal ECT; BL = Bilateral rTMS; BF = Bifrontal ECT; tDCS = transcranial Direct Current Stimulation; sTMS = synchronised Transcranial Magnetic Stimulation; pTMS = priming Transcranial Magnetic Stimulation; TBS = Theta Burst Stimulation; iTBS = intermittent TBS; dTMS = deep Transcranial Magnetic Stimulation; cTBS = continuous TBS; blTBS = Bilateral TBS; aTMS = accelerated Transcranial Magnetic Stimulation.

Comparing active treatments, BT ECT was associated with higher response than BF ECT, LM-RUL ECT, LF-L rTMS, cTBS and dTMS. H-RUL ECT was associated with higher response than LM-RUL ECT and cTBS. pTMS and BL rTMS were more efficacious than cTBS. No other significant differences between active treatments were found (Table 1).

All-cause discontinuation rates were available for 227 treatment arms (n = 6,362), including all 18 active interventions and sham (Figure 4).

**Figure 4.**
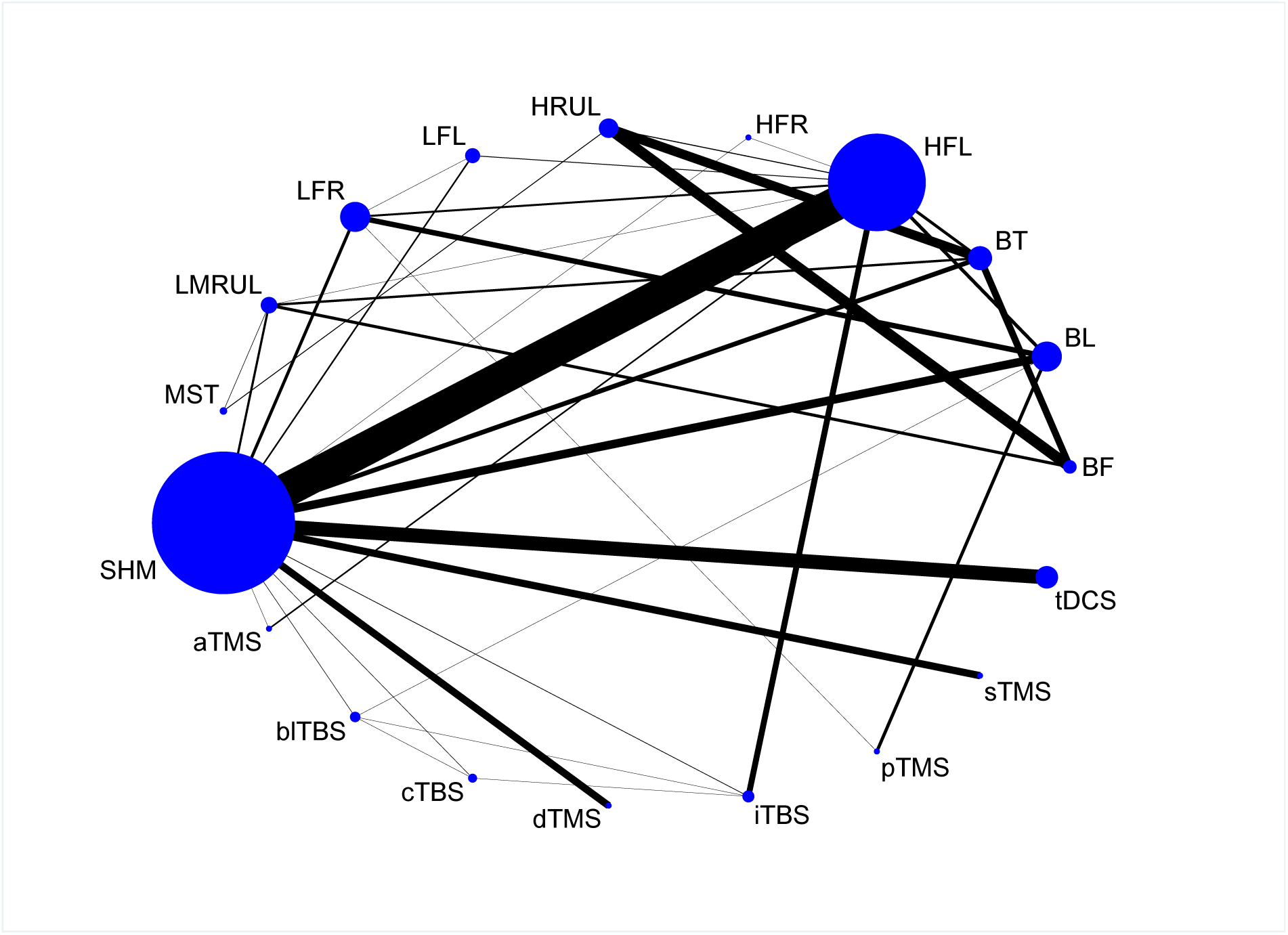
Network plot of available treatment comparisons for all-cause discontinuation rates. The size of the nodes is proportional to the number of patients randomised to each treatment. The width of the lines is proportional to the number of RCTs comparing each pair of treatments. SHM = Sham; MST = Magnetic Seizure Therapy; ECT = Electroconvulsive Therapy; LMRUL = Low to Moderate-Dose Right Unilateral ECT; rTMS = repetitive Transcranial Magnetic Stimulation; LFR = Low-Frequency Right rTMS; LFL = Low-Frequency Left rTMS; HRUL = High-Dose Right Unilateral ECT; HFR = High-Frequency Right rTMS; HFL = High-Frequency Left rTMS; BT = Bitemporal ECT; BL = Bilateral rTMS; BF = Bifrontal ECT; tDCS = transcranial Direct Current Stimulation; sTMS = synchronised Transcranial Magnetic Stimulation; pTMS = priming Transcranial Magnetic Stimulation; TBS = Theta Burst Stimulation; iTBS = intermittent TBS; dTMS = deep Transcranial Magnetic Stimulation; cTBS = continuous TBS; blTBS = Bilateral TBS; aTMS = accelerated Transcranial Magnetic Stimulation.

The NMA results suggest that pTMS was more acceptable than LF-L rTMS (OR=0.11, 95%CI 0.02-0.59), MST (0.13, 0.02-0.95), aTMS (0.16, 0.03-0.93), tDCS (0.18, 0.05-0.61), LF-R rTMS (0.23, 0.08-0.72), dTMS (0.25, 0.07-0.92), HF-L rTMS (0.26, 0.08-0.79) and sham (0.21, 0.07-0.65). Moreover, BL rTMS was associated with fewer drop-outs than tDCS and sham (Table 1). All treatments were at least as acceptable as sham and these conclusions did not change when examining prediction intervals (Figure 5).

**Figure 5.**
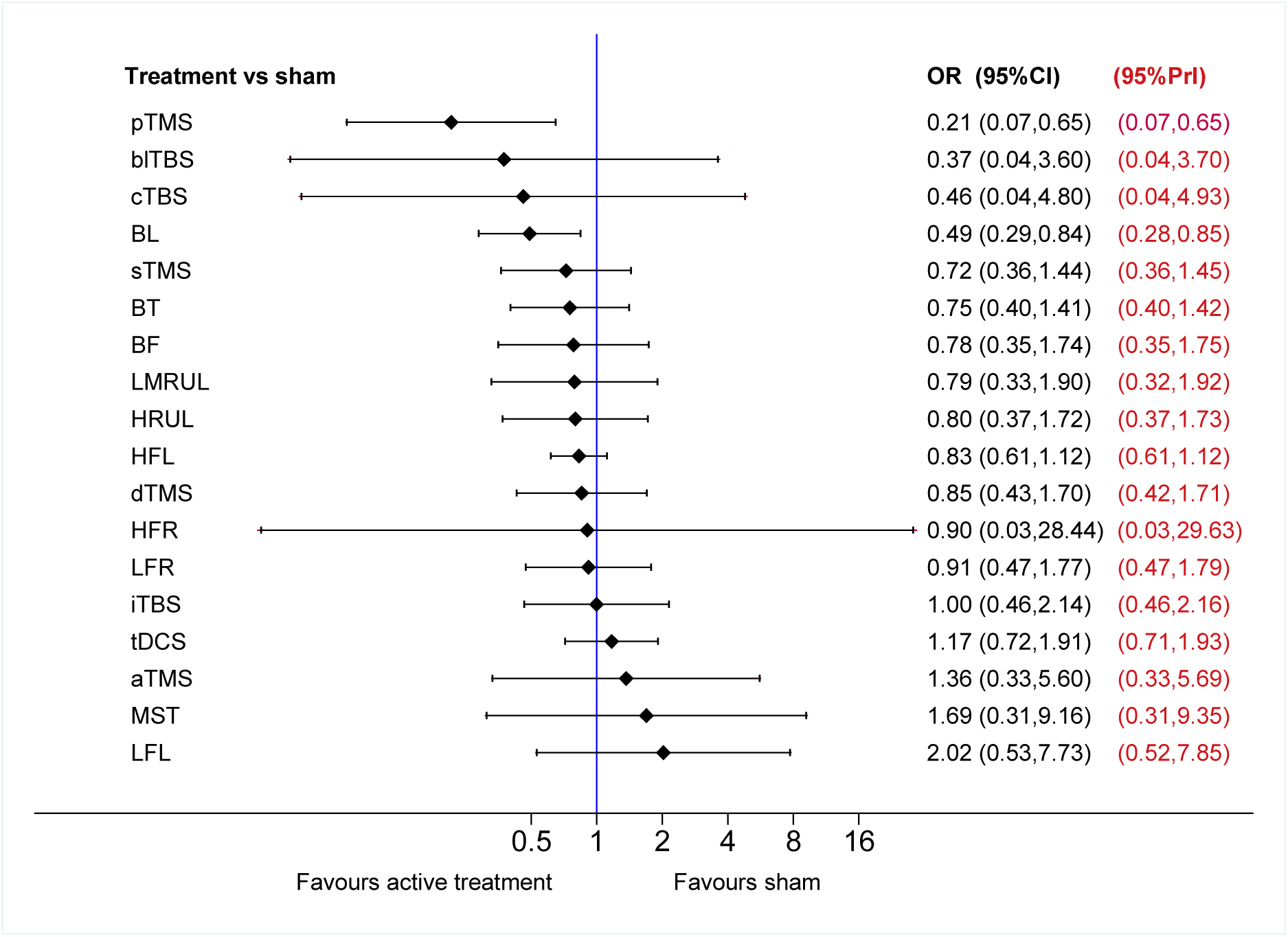
Forest plot of active vs sham treatment comparisons for all-cause discontinuation rates. Effect sizes represent relative odds ratios (ORs) with 95% confidence intervals (Cis) and 95% prediction intervals (PrIs). SHM = Sham; MST = Magnetic Seizure Therapy; ECT = Electroconvulsive Therapy; LMRUL = Low to Moderate-Dose Right Unilateral ECT; rTMS = repetitive Transcranial Magnetic Stimulation; LFR = Low-Frequency Right rTMS; LFL = Low-Frequency Left rTMS; HRUL = High-Dose Right Unilateral ECT; HFR = High-Frequency Right rTMS; HFL = High-Frequency Left rTMS; BT = Bitemporal ECT; BL = Bilateral rTMS; BF = Bifrontal ECT; tDCS = transcranial Direct Current Stimulation; sTMS = synchronised Transcranial Magnetic Stimulation; pTMS = priming Transcranial Magnetic Stimulation; TBS = Theta Burst Stimulation; iTBS = intermittent TBS; dTMS = deep Transcranial Magnetic Stimulation; cTBS = continuous TBS; blTBS = Bilateral TBS; aTMS = accelerated Transcranial Magnetic Stimulation.

Findings pertaining to the secondary and tertiary efficacy measures (remission and continuous post-treatment depression severity scores) are shown in Supplement 8-10.

#### Ranking probabilities

Ranking plots for all outcomes are presented in Supplement 11. The most efficacious treatments in terms of response were BT ECT (35.6%) and pTMS (19.3%), while LF-L rTMS (30.3%) and cTBS (29.7) were least efficacious. In terms of all-cause discontinuation, pTMS (40.6%) and blTBS (22.8%) had the highest probabilities of being best accepted while LF-L rTMS (28.2%) and HF-R rTMS (23.4%) had similar probabilities of being least accepted.

#### Inconsistency

Fitting the design-by-treatment interaction model provided no evidence for significant inconsistency for response, remission and all-cause discontinuation (global Wald tests: *p* =0.42-0.99). However, there was some evidence for inconsistency in the post-treatment depression severity network (global Wald test: *p* = 0.09). We present inconsistency plots for each outcome in Supplement 12. For our primary outcome measure of efficacy (response), we found evidence for inconsistency in 3/21 (14%) loops, while there was no evidence for inconsistency for all-cause discontinuation.

### Sensitivity analysis

Excluding trials that investigated tDCS did not materially change our results and overall conclusions (Supplement 13). When trials with high overall risk of bias were excluded, MST and iTBS were no longer associated with higher response than sham. There was also no evidence that pTMS was associated with fewer drop-outs than any other treatment in the network.

## Discussion

This is the most comprehensive systematic review and network meta-analysis of non-surgical brain stimulation for the acute treatment of major depressive episodes in adults. We included data from 113 clinical trials including 6,750 patients with MDD or bipolar depression who were randomised to 18 distinct treatment protocols or sham. The quality of the evidence was typically of low or unclear risk of bias (92 out of 113 trials; 81.4%).

Our findings provide evidence for the antidepressant efficacy of ECT. Previous comparative analyses did not consistently favour BT ECT or RUL ECT, and it has been suggested that RUL ECT needs to be delivered at multiples of seizure threshold to be effective^31,32^. Trials that employed electrical dosages at or just above seizure threshold may have underestimated treatment effects. Our findings support this view. We found no evidence of differences in efficacy between H-RUL ECT and BT ECT across outcomes, while LM-RUL ECT (i) was less efficacious than BT ECT across outcomes in pairwise meta-analyses, (ii) was associated with lower response rates than BT ECT and H-RUL ECT in NMA and (iii) failed to separate from sham.

Two trials^33,34^ evaluated the antidepressant efficacy of MST compared to moderate-dose RUL ECT and one trial^35^ compared MST to H-RUL ECT. While we found no evidence of differences between treatments in pairwise meta-analysis, the NMA of response provides preliminary evidence in favour of MST compared to sham. However, this estimate relies on indirect evidence only and a sham-controlled trial is needed to confirm this finding.

Consistent with previous analyses^14,36-39^ our results provide evidence for the antidepressant efficacy of HF-L and LF-R rTMS. The efficacy of BL rTMS is comparable to both HF-L and LF-R rTMS^11^, with little evidence for additional benefit of bilateral compared to unilateral stimulation. The finding that neither LF-L nor HF-R rTMS were more efficacious than sham lends support to the view that the antidepressant effects of rTMS depend on specific stimulation frequency and coil location.

We found limited evidence in support of the more recent treatment modalities. Compared to sham, iTBS and pTMS were associated with improved response and remission in NMA, while blTBS was associated with higher response. However, when considering data from pairwise direct comparisons only, the evidence in favour of iTBS compared to sham was limited to higher response. With respect to dTMS we found evidence of antidepressant efficacy across outcome measures in pairwise analyses but not in NMA. Considering that the direct evidence is based on data from two RCTs^40,41^ only, further investigations are warranted. We found no evidence suggesting that cTBS, aTMS and sTMS are effective treatments for major depressive episodes. However, these findings need to be treated with caution due to the limited number of included studies. Finally, while previous meta-analyses of the antidepressant efficacy of tDCS yielded inconsistent results^10,42-46^, we found tDCS to be efficacious across outcomes in both pairwise and network meta-analyses.

There was little evidence for differences in all-cause discontinuation between active treatments and sham. The notable exception was pTMS for which lower drop-out rates were reported. However, we did not examine specific undesired and adverse effects associated with treatment.

Limitations were that most included studies exhibited unclear risk of bias, particularly with respect to random sequence generation and allocation concealment. Overall risk of bias was deemed high in 21 trials (18.6%). In a sensitivity analysis excluding these trials we found that iTBS and MST were no longer associated with higher response than sham. Moreover, we found no evidence of differences in all-cause discontinuation between pTMS and other treatments.

There was some evidence for statistical heterogeneity within pairwise comparisons and a small number of loops in our NMA of response suggested inconsistency between direct and indirect sources of evidence. To facilitate interpretation of our results taking the magnitude of heterogeneity into account, we presented predictive intervals for all sham-comparisons. For MST, iTBS and tDCS the estimate of a future trial might suggest that these treatment protocols are no more efficacious than sham.

While several RCTs have compared different rTMS or different ECT protocols, few trials have compared novel brain stimulation techniques to ECT. A conceivable explanation is that rTMS and related interventions require no anaesthetic but a higher level of cooperation from the patient, whereas ECT can be prescribed to patients who are more severely depressed. However, most trials that were included in our analyses were conducted after multiple pharmacotherapies had failed and patient characteristics did not materially differ between most treatment comparisons. Trials that examined tDCS were excluded in a sensitivity analysis because these studies showed some differences with other treatment comparisons and because tDCS is a less invasive treatment protocol. Excluding these studies did not materially change our results.

Finally, we focused on the acute antidepressant effects at primary study endpoint and our conclusions might not apply to the long-term effects of non-surgical brain stimulation. Continuation and maintenance treatment will need to be reviewed separately.

Our findings have implications for clinical decision-making and research. They inform clinicians, patients and healthcare providers on the comparative efficacy and acceptability of multiple non-surgical brain stimulation techniques. Moreover, they are relevant to policy makers involved in regulating medical devices and developing treatment guidelines. This review also highlights important research priorities in the field of brain stimulation, for instance the need to conduct further well-designed RCTs comparing novel treatment modalities and sham-controlled trials investigating MST.

## Conclusion

We found that non-surgical brain stimulation techniques constitute viable alternative or add-on treatments for adult patients with major depressive episodes. Our findings also highlight the need to consider other patient and treatment-related factors in addition to antidepressant efficacy and acceptability when making clinical decisions.

## Authorship contributions

JM conceived and supervised the study; JM and VV independently performed the literature search and conducted the risk of bias assessment; JM extracted, analysed and interpreted the data; VV independently reviewed the extracted data; JM wrote the paper with input from VV, BC, RH, CHYF and AHY. All authors read and approved the final version of the paper.

## Acknowledgments

A preliminary version of this work was performed as partial fulfilment towards the International Master in Affective Neuroscience of Maastricht University and the University of Florence.

## Supplementary material

Supplementary information is available online.

## References

1 Kessler, R. C. et al. The epidemiology of major depressive disorder: results from the National Comorbidity Survey Replication (NCS-R). JAMA 289, 3095–3105 (2003).

2 Murray, C. J. et al. Global, regional, and national disability-adjusted life years (DALYs) for 306 diseases and injuries and healthy life expectancy (HALE) for 188 countries, 1990–2013: quantifying the epidemiological transition. The Lancet 386, 2145–2191 (2015).

3 Fekadu, A. et al. What happens to patients with treatment-resistant depression? A systematic review of medium to long term outcome studies. Journal of Affective Disorders 116, 4–11 (2009).

4 Rush, A. J. et al. Acute and longer-term outcomes in depressed outpatients requiring one or several treatment steps: a STAR* D report. American Journal of Psychiatry 163, 1905–1917 (2006).

5 Kikuchi, T., Suzuki, T., Uchida, H., Watanabe, K. & Mimura, M. Association between antidepressant side effects and functional impairment in patients with major depressive disorders. Psychiatry Research 210, 127–133 (2013).

6 Serretti, A. & Mandelli, L. Antidepressants and body weight: a comprehensive review and meta-analysis. Journal of Clinical Psychiatry 71, 1259–1272 (2010).

7 Papakostas, G. I. Tolerability of modern antidepressants. The Journal of Clinical Psychiatry 69, 8–13 (2008).

8 Velligan, D. I. et al. The expert consensus guideline series: adherence problems in patients with serious and persistent mental illness. Journal of Clinical Psychiatry 70, 1–46 (2009).

9 Scott, J. & Young, A. H. Psychotherapies should be assessed for both benefit and harm. The British Journal of Psychiatry 208, 208–209 (2016).

10 Mutz, J., Edgcumbe, D. R., Brunoni, A. R. & Fu, C. H. Efficacy and acceptability of non-invasive brain stimulation for the treatment of adult unipolar and bipolar depression: a systematic review and meta-analysis of randomised sham-controlled trials. Neuroscience & Biobehavioral Reviews (2018).

11 Chen, J.-j. et al. Bilateral vs. unilateral repetitive transcranial magnetic stimulation in treating major depression: a meta-analysis of randomized controlled trials. Psychiatry Research 219, 51–57 (2014).

12 Salanti, G. Indirect and mixed-treatment comparison, network, or multiple-treatments meta-analysis: many names, many benefits, many concerns for the next generation evidence synthesis tool. Research Synthesis Methods 3, 80–97 (2012).

13 Leucht, S. et al. Network meta-analyses should be the highest level of evidence in treatment guidelines. European Archives of Psychiatry and Clinical Neuroscience 6, 477–480 (2016).

14 Brunoni, A. R. et al. Repetitive transcranial magnetic stimulation for the acute treatment of major depressive episodes: a systematic review with network meta-analysis. JAMA Psychiatry 74, 143–152 (2017).

15 Chen, J.-j., Zhao, L.-b., Liu, Y.-y., Fan, S.-h. & Xie, P. Comparative efficacy and acceptability of electroconvulsive therapy versus repetitive transcranial magnetic stimulation for major depression: a systematic review and multiple-treatments meta-analysis. Behavioural Brain Research, 30–36 (2017).

16 Hutton, B. et al. The PRISMA extension statement for reporting of systematic reviews incorporating network meta-analyses of health care interventions: checklist and explanations. Annals of Internal Medicine 162, 777–784 (2015).

17 Hamilton, M. A rating scale for depression. Journal of Neurology, Neurosurgery & Psychiatry 23, 56–62 (1960).

18 Montgomery, S. A., & Åsberg, M. A new depression scale designed to be sensitive to change. The British Journal of Psychiatry 134, 382–389 (1979).

19 Higgins, J. P. & Green, S. Cochrane Handbook for Systematic Reviews of Interventions Version 5.1.0. The Cochrane Collaboration, 33–49 (2011).

20 Higgins, J. P. et al. The Cochrane Collaboration’s tool for assessing risk of bias in randomised trials. BMJ 343, 889–893 (2011).

21 Schwarzer, G. Meta: An R package for meta-analysis. R News 7, 40–45 (2007).

22 Higgins, J. P., Thompson, S. G., Deeks, J. J. & Altman, D. G. Measuring inconsistency in meta-analyses. BMJ 327, 557–561 (2003).

23 Chaimani, A., Higgins, J. P., Mavridis, D., Spyridonos, P. & Salanti, G. Graphical tools for network meta-analysis in STATA. PloS One 8, e76654 (2013).

24 Higgins, J. et al. Consistency and inconsistency in network meta-analysis: concepts and models for multi-arm studies. Research Synthesis Methods 3, 98–110 (2012).

25 White, I. R., Barrett, J. K., Jackson, D. & Higgins, J. P. Consistency and inconsistency in network meta-analysis: model estimation using multivariate meta-regression. Research Synthesis Methods 3, 111–125 (2012).

26 White, I. R. Multivariate random-effects meta-analysis. Stata Journal 9, 40 (2009).

27 White, I. R. Multivariate random-effects meta-regression: updates to mvmeta. Stata Journal 11, 255 (2011).

28 White, I. R. Network meta-analysis. Stata Journal 15, 951–985 (2015).

29 Fitzgerald, P. et al. A randomized trial of unilateral and bilateral prefrontal cortex transcranial magnetic stimulation in treatment-resistant major depression. Psychological Medicine 41, 1187–1196 (2011).

30 Pallanti, S., Bernardi, S., Di Rollo, A., Antonini, S. & Quercioli, L. Unilateral low frequency versus sequential bilateral repetitive transcranial magnetic stimulation: is simpler better for treatment of resistant depression? Neuroscience 167, 323–328 (2010).

31 Sackeim, H. A. et al. Effects of stimulus intensity and electrode placement on the efficacy and cognitive effects of electroconvulsive therapy. New England Journal of Medicine 328, 839–846 (1993).

32 McCall, W. V., Reboussin, D. M., Weiner, R. D. & Sackeim, H. A. Titrated moderately suprathreshold vs fixed high-dose right unilateral electroconvulsive therapy: acute antidepressant and cognitive effects. Archives of General Psychiatry 57, 438–444 (2000).

33 Fitzgerald, P. B. et al. A pilot study of the comparative efficacy of 100 Hz magnetic seizure therapy and electroconvulsive therapy in persistent depression. Depression and Anxiety (2018).

34 Kayser, S. et al. Antidepressant effects, of magnetic seizure therapy and electroconvulsive therapy, in treatment-resistant depression. Journal of Psychiatric Research 45, 569–576 (2011).

35 Kayser, S. et al. Degree of postictal suppression depends on seizure induction time in magnetic seizure therapy and electroconvulsive therapy. The Journal of ECT 33, 167–175 (2017).

36 Schutter, D. Antidepressant efficacy of high-frequency transcranial magnetic stimulation over the left dorsolateral prefrontal cortex in double-blind sham-controlled designs: a meta-analysis. Psychological Medicine 39, 65–75 (2009).

37 Berlim, M. T., Van den Eynde, F., Tovar-Perdomo, S. & Daskalakis, Z. Response, remission and drop-out rates following high-frequency repetitive transcranial magnetic stimulation (rTMS) for treating major depression: a systematic review and meta-analysis of randomized, double-blind and sham-controlled trials. Psychological Medicine 44, 225–239 (2014).

38 Lepping, P. et al. A systematic review of the clinical relevance of repetitive transcranial magnetic stimulation. Acta Psychiatrica Scandinavica 130, 326–341 (2014).

39 Berlim, M. T., Van den Eynde, F. & Daskalakis, Z. J. Clinically meaningful efficacy and acceptability of low-frequency repetitive transcranial magnetic stimulation (rTMS) for treating primary major depression: a meta-analysis of randomized, double-blind and sham-controlled trials. Neuropsychopharmacology 38, 543–551 (2013).

40 Levkovitz, Y. et al. Efficacy and safety of deep transcranial magnetic stimulation for major depression: a prospective multicenter randomized controlled trial. World Psychiatry 14, 64–73 (2015).

41 Tavares, D. F. et al. Treatment of bipolar depression with deep TMS: results from a double-blind, randomized, parallel group, sham-controlled clinical trial. Neuropsychopharmacology (2017).

42 Berlim, M. T., Van den Eynde, F. & Daskalakis, Z. J. Clinical utility of transcranial direct current stimulation (tDCS) for treating major depression: a systematic review and meta-analysis of randomized, double-blind and sham-controlled trials. Journal of Psychiatric Research 47, 1–7 (2013).

43 Kalu, U., Sexton, C., Loo, C. & Ebmeier, K. Transcranial direct current stimulation in the treatment of major depression: a meta-analysis. Psychological Medicine 42, 1791–1800 (2012).

44 Shiozawa, P. et al. Transcranial direct current stimulation for major depression: an updated systematic review and meta-analysis. International Journal of Neuropsychopharmacology 17, 1443–1452 (2014).

45 Meron, D., Hedger, N., Garner, M. & Baldwin, D. S. Transcranial direct current stimulation (tDCS) in the treatment of depression: systematic review and meta-analysis of efficacy and tolerability. Neuroscience & Biobehavioral Reviews 57, 46–62 (2015).

46 Brunoni, A. R. et al. Transcranial direct current stimulation for acute major depressive episodes: meta-analysis of individual patient data. The British Journal of Psychiatry, doi:10.1192/bjp.bp.115.164715 (2016).

